# Next-generation sequencing of pIJ101-based *Streptomyces-E. coli* shuttle vectors

**DOI:** 10.1101/2020.02.26.966358

**Authors:** Katelyn Brown, S. Eric Nybo

## Abstract

The sequences of two commonly used pIJ101-based *Streptomyces-E. coli* shuttle vectors, pEM4 and pUWL201, were determined using next-generation sequencing and mapped. pEM4 was found to be 8.3 kbp long, containing genes encoding β-lactamase, thiostrepton resistance, the *lacZ*□ fragment, an *E. coli* pUC19 origin of replication, and the *Streptomyces* pIJ101 origin of replication. pUWL201 was found to be 6.78 kbp long, containing genes encoding β-lactamase, thiostrepton resistance, the *lacZ*□ fragment, an *E. coli* pUC19 origin of replication, and the *Streptomyces* pIJ101 origin of replication. Interestingly, the sequences for both pEM4 and pUWL201 exceed their previously reported size by 1.1 kbp and 0.4 kbp, respectively. This report updates the literature with the corrected sequences for these two popular vectors, ensuring their compatibility with modern synthetic biology cloning methodologies.

## Introduction

Next-generation sequencing (NGS) techniques, such as RNA-seq and Illumina whole genome sequencing (WGS), have transformed the biological sciences via deep-sequencing of whole-cell transcriptomic profiles and genomes at greatly reduced cost (Li et al., 2010; Wang et al., 2009)WGS relies on deep-sequencing to generate many smaller reads that are then aligned to a reference file. In the absence of a reference file, de novo sequencing of genomes relies on the generation of millions of parallel reads that are then compiled into contigs and assembled based on the depth of coverage afforded by NGS platforms (Li et al., 2010). De novo sequencing can also be applied to the determination of unknown plasmid sequences with great accuracy and has supplanted Sanger sequencing for in some laboratory synthetic biology workflows (Suckling et al., 2019). De novo sequencing is a powerful tool for the determination of plasmid vector sequences for plasmids that have widespread use for several decades, but for which no sequence information is available in the literature.

pIJ101 is a high-copy plasmid vector that is replicated in *Streptomyces lividans* and *Streptomyces coelicolor* at a copy number of approximately 70-100 copies per cell (Kieser et al., 1982). Since the first report of its use in cloning of *Streptomyces* genes in the 1980s, many different derivatives of pIJ101 have been developed, including pWHM3 and pUWL201 (Doumith et al., 2000; Vara et al., 1989). In early reports, cloning via restriction enzymes into sites in a multiple-cloning site (MCS) was the gold-standard technique for studying gene expression in *Streptomyces*, therefore, there was little interest in publishing the complete nucleotide sequence of the various reported pIJ101 derivatives. However, modern cloning techniques are sequence-specific, including Golden-gate assembly (Engler et al., 2008), Gibson assembly (Gibson et al., 2009), and even yeast recombination assembly (Joska et al., 2014), methods for which these plasmid vectors would be incompatible without exact sequencing information. Therefore, in this report, we determined the sequence for the popular pEM4 and pUWL201 expression vectors using next-generation sequencing.

### HIGHLIGHTS

Two closely-related *E. coli*-*Streptomyces* shuttle vectors were sequenced and analyzed. Updated sequences and vector maps for pEM4 and pUWL201 facilitate incorporation of these plasmids into modern synthetic biology workflows.

## Materials and Methods

### Bacterial strains and growth conditions

*Escherichia coli* JM109 (Promega) was used to propagate plasmids for sequencing analysis. *E. coli* JM109 was grown lytic broth medium, and plasmids were introduced into *E. coli* by chemically competent transformation done by standard procedures (Sambrook and W Russell, 2001). For *E. coli* strains harboring plasmids, ampicillin was added at a final concentration of 100 µg/mL. Plasmid pEM4 was provided by Dr. Jurgen Rohr (University of Kentucky) and pUWL201 was provided by Dr. Steven van Lanen (University of Kentucky.

### Next-generation plasmid sequencing

Whole plasmid verification was carried out at the Massachusetts General Hospital Center for Computational and Integrative Biology DNA Core. Next-generation sequencing was carried out on an Illumina MiSeq platform with V2 chemistry. *De novo* plasmid sequence assembly was achieved via the UltraCycler v1.0 assembler (Brian Seed and Huajun Wang, unpublished).

## Results and Discussion

### Sequencing and analysis of pEM4 and pUWL201

Plasmids pEM4 and pUWL201 were sequenced via the Illumina MiSeq next-generation sequencing platform. pEM4 is a high-copy number, *E. coli*-*Streptomyces spp*. shuttle vector which is derived from pWHM4 *via* insertion of the strong *ermE*p* promoter from *Saccharopolyspora erythraea* (Quirós et al., 1998). The *ermE*p* promoter (i.e. the “*ermE-*up” promoter) is a mutated version of the *ermE* promoter that features a TGG deletion of the *ermE* P1 promoter (Bibb et al., 1986; Schmitt-John and Engels, 1992). pEM4 (GenBank Accession No. MN970094) was determined to be an 8,289 base pair plasmid, which features an additional 1,000 base pairs of sequence as compared to the originally reported plasmid size of 7,200 base pairs (Vara et al., 1989). The extra base pairs of sequence include 300 base pairs for cloning of the *ermE*p* promoter and 700 base pairs from pIJ486 encoding a partial ATP-binding cassette transporter carried over from the original cloning scheme. pEM4 contains the *bla* gene encoding ampicillin resistance, the *tsr* 23S-rRNA methyltransferase from *Streptomyces azureus* encoding thiostrepton resistance, the high-copy number pMB1 origin of replication from pUC19, and the pIJ702 origin of replication for *Streptomyces* spp. (Table 1 and Figure 1).

**Table 1.** Plasmid features of pEM4 and pUWL201.

| **Plasmid Features** | **pEM4** | **pUWL201** |
| --- | --- | --- |
| Size (bp) | 8289 bp | 6780 bp |
| G + C content (%) | 62.6% | 57.8% |
| ColE1 origin | 6901...7583 | 6383...285 |
| pIJ702 origin | 3847...5433 | — |
| pIJ101 origin | — | 1052...2422 |
| bla resistance | 7678...249 | 5626...6285 |
| tsr resistance | 1607...2416 | 3564...4373 |
| ermE*p promoter | 892...933 | 633...671 |
| Polylinker | 846...1073 | 509...632 |
| LacZ alpha | 688...756 | 4772...4840 |

— Feature is not present in the vector.

**Figure 1.**
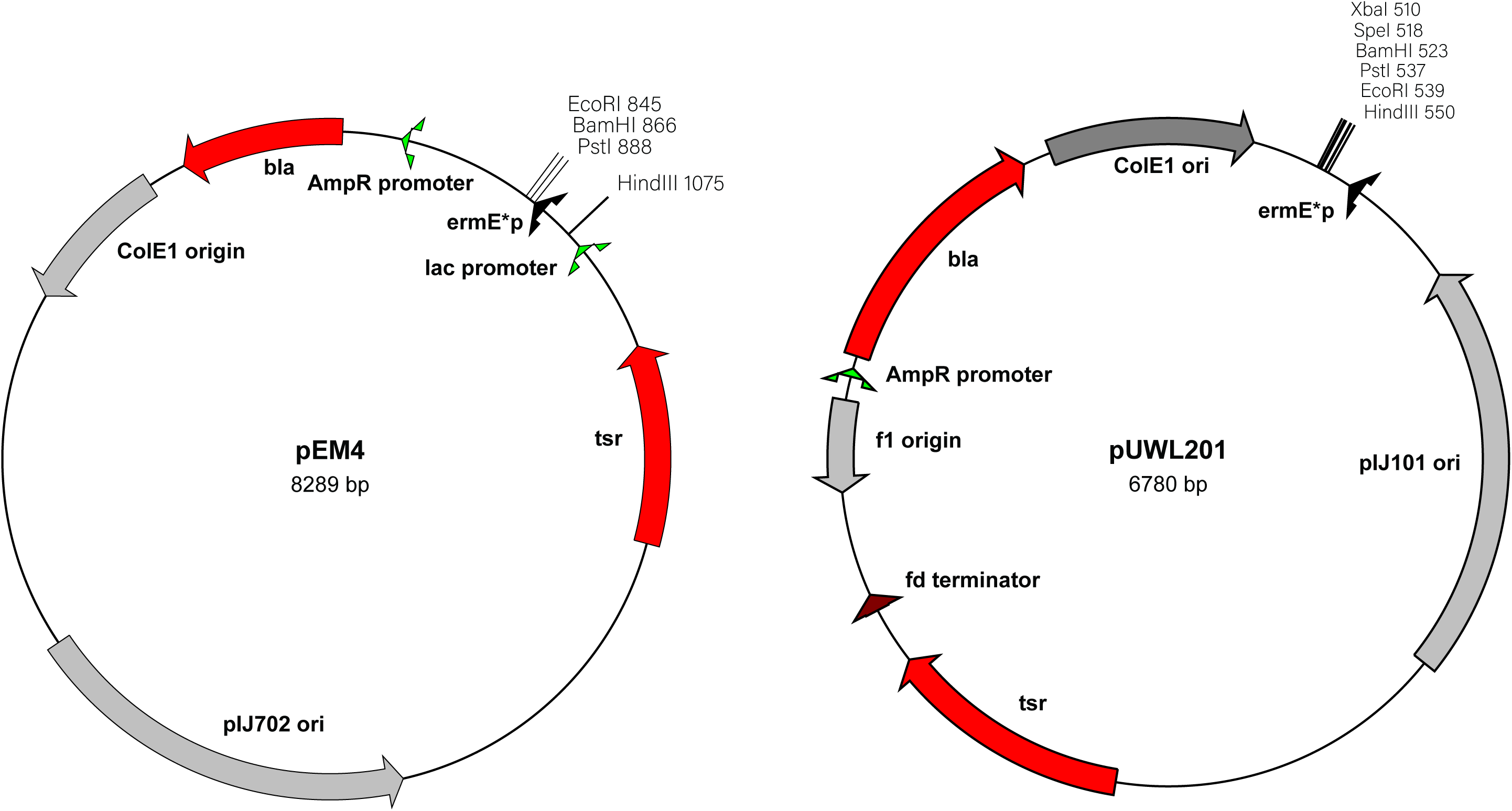
Plasmids maps of pEM4 and pUWL201.

pUWL201 (GenBank Accession No. MN992950) is derived from pIJ4070 via introduction of a 280 base pair fragment encoding the *ermE*p* promoter into the *Kpn*I – *Xba*I sites of the polylinker region (Doumith et al., 2000). pUWL201 was determined to be 6,780 base pairs in size, which is 380 base pairs larger than the originally reported plasmid size of 6,400 base pairs. Sequencing of the multiple cloning site revealed the presence of an additional *Xba*I restriction site. pUWL201 was determined to have the *bla* gene encoding ampicillin resistance, the *tsr* 23S-rRNA methyltransferase from *S. azureus* encoding thiostrepton resistance, the ColE1 origin of replication for *E. coli*, and the pIJ101 origin of replication for *Streptomyces* spp. (Table 1 and Figure 1).

## Conclusion

The sequences for the broadly used pEM4 and pUWL201 *E. coli*-*Streptomyces* shuttle vectors were determined via next-generation sequencing and analyzed. Both vectors revealed additional nucleotide sequences and were larger in size than previously reported. The characterization of these vectors reveals that vectors traditionally used for genetic engineering of prokaryotes may contain carryover sequence not explicitly required for plasmid function. This work should facilitate efforts to generate “minimal” versions of these *E. coli – Streptomyces* spp. vectors and will enable their adaptation for synthetic biology.

## Acknowledgements

Dr. Jurgen Rohr and Dr. Steven Van Lanen (University of Kentucky) are gratefully acknowledged for the gift of plasmids pEM4 and pUWL201. This study was funded by start-up funds from Ferris State University to S.E.N.

